# The protein interactome of the citrus Huanglongbing pathogen *Candidatus* Liberibacter asiaticus

**DOI:** 10.1101/2023.07.10.548374

**Authors:** Erica W. Carter, Orlene Guerra Peraza, Nian Wang

## Abstract

*Candidatus* Liberibacter asiaticus (CLas) is the causal agent of the devastating citrus Huanglongbing (HLB) disease. Our understanding of the pathogenicity mechanism and biology of CLas remain limited because CLas has not been cultured in artificial media. CLas encodes 1136 proteins of which 415 have unknown functions. Since genetic studies of CLas genes with unknown functions are impossible, we utilized genome-wide protein-protein interactions (PPIs) yeast-two-hybrid (Y2H) assays to help solve the mystery. PPIs are fundamental to all cellular processes and machinery and instrumental in investigating uncharacterized proteins and inferring biological pathways. In total, 916 bait and 936 prey proteins were included in the three-phase screening, which identified 4245 interactions. The false positive rate of the Y2H assay was estimated to be 3.1%. Pull-down assays confirmed the robustness of our Y2H. The average interactions per node for CLas Y2H interactome were approximately 15.6, significantly higher than free-living bacteria, indicating genome reduction has led to a multi-function of proteins. PPIs provide clues for functions of 371 uncharacterized proteins of CLas. Forty HUB node proteins were identified which might play critical roles in CLas, including a quinone oxidoreductase and LysR that are known to protect bacteria against oxidative stress. This explains why CLas survives well in the phloem even though it triggers immune-mediated disease, systemic and chronic production of reactive oxygen species, and phloem cell death. This PPI database facilitates the investigation of CLas cellular biochemistry and physiology, functions of uncharacterized proteins, and pathogenicity mechanisms of the pathogen.

## Main

Citrus Huanglongbing (HLB) is one of the most destructive plant diseases worldwide, and no effective and economic disease control measures are available for HLB-endemic citrus production regions^1^. HLB has devastated the citrus industry in many regions, including Florida, US^2^. HLB is caused by *Candidatus* Liberibacter asiaticus (CLas), *Ca*. L. americanus, and *Ca*. L. africanus, with CLas being the most prevalent. *Ca*. Liberibacter species are endophytic bacteria that colonize the phloem tissues of citrus plants and are transmitted by the hemipteran insect *Diaphorina citri*^3, 4^ The environments inside the phloem tissue and psyllids are nutrient rich. Consequently, *Ca*. Liberibacter species have undergone reductive evolution and have considerably smaller genomes (approximately 1.2 MB) than their culturable, free-living relatives, such as *Agrobacterium*^3, 4^. Cultivation of HLB pathogens in artificial media has not been achieved despite extensive effort^5^, which prevented genetic manipulation and impeded understanding of their biology and pathogenicity mechanism.

The CLas genome encodes approximately 1,027 proteins, of which 612 have predicated functions based on sequence similarities with characterized proteins from culturable microorganisms. However, 415 CLas proteins have no known functions^4^. Investigating protein functions, particularly those with unknown roles, is critical for elucidating the biology and pathogenicity of CLas and developing effective strategies to combat this notorious citrus disease. Protein–protein interactions (PPIs) are fundamental to all cellular processes and machinery. PPIs modify enzyme and protein activities, catalyze metabolic reactions, and activate signaling pathways^6^. Interactions with proteins of known functions provide informative clues for function prediction of uncharacterized proteins^7^. Exploring PPIs was also used to investigate biological pathways^8^. Furthermore, targeting PPIs can be used to develop novel antimicrobials^9^. The Yeast Two-hybrid system (Y2H) is the most commonly used approach to investigate interactomes and has been elegantly applied to various organisms, such as animals, *Drosophila melanogaster*^10^ and *Caenorhabditis elegans*^11, 12^; plants, *Arabidopsis thaliana*^13, 14^; fungi, *Saccharomyces cerevisiae*^7^; and bacteria, *Streptococcus pneumoniae*^15^, *Treponema pallidum*^16^, *Campylobacter jejuni*^17^, *Synechocystis sp.*^18^, *Mycobacterium tuberculosis*^19^, *Mesorhizobium loti*^20^, *Escherichia coli*^21^, *Bacillus subtilis*^22^, and *Helicobacter pylori*^8, 23^. Affinity Purification Mass Spectrometry (AP-MS) has also been used to study the *Mycoplasma pneumoniae* interactome^6, 24^ (Extended Data Table S1). However, AP-MS detects PPIs in the native organism, which prevents its application in non-cultured microorganisms such as CLas, whereas Y2H, using a non-native host: yeast, bypasses this limitation. Interactome studies (Extended Data Table S1) have led to significant progress in inferring the function of unknown proteins, elucidating protein organization of a cell, identifying novel functions of known proteins, and in some cases, are used to find interaction points between pathogenic organisms and their eukaryotic hosts ^25–31^. Here, we conducted a genome-wide Y2H interactome study of CLas.

### High-throughput Y2H screening

We conducted Y2H assays to identify the CLas interactome. Full-length open reading frames (ORFs) were used for yeast vector construction to increase the possibility of identifying biologically relevant interactions. 974 ORFs were PCR amplified (Supplementary Dataset S1), and 942 ORFs were successfully cloned (Extended Data Figure S1A). In total, 926 “bait” proteins in pGEO_BD, which was modified from pGBKT7_BD (Extended Figure S1B), and 950 “prey” proteins in pGAD_T7 remained. After the removal of autoactivating constructs (Extended Data Figure S2), 916 “bait” and 936 “prey” constructs were used in the screening, representing 91% of genes encoded by CLas. We constructed the CLas binary interactome using a three-phase array screening method (Figure 1A). The first two pooled array phases removed non-interactors. Proteins interacting on the most stringent media, QX (-trp, -leu, -ade, -his, +XαGal) and QXA (-trp, -leu, -ade, -his, +XαGal, +AbA) in the first two screening phases were used in the pairwise array in the third screening phase. The pairwise screening enabled us to identify all potential interactions from the mini-pooled arrays and detect the interacting partners. In addition, pairwise rescreening eliminated probable promiscuous proteins from the dataset. The final (third phase) screening between 617 baits and 466 preys resulted in 4245 interactions between 542 proteins (Supplementary Dataset S2), covering 52.8% of CLas genes, denoted as the CLas_whole network (Figure 1B). Among the 542 proteins, 371 had known functions, whereas 171 were uncharacterized proteins (Supplementary Dataset S3; Extended Data Tables S2 and S3).

**Figure 1.**
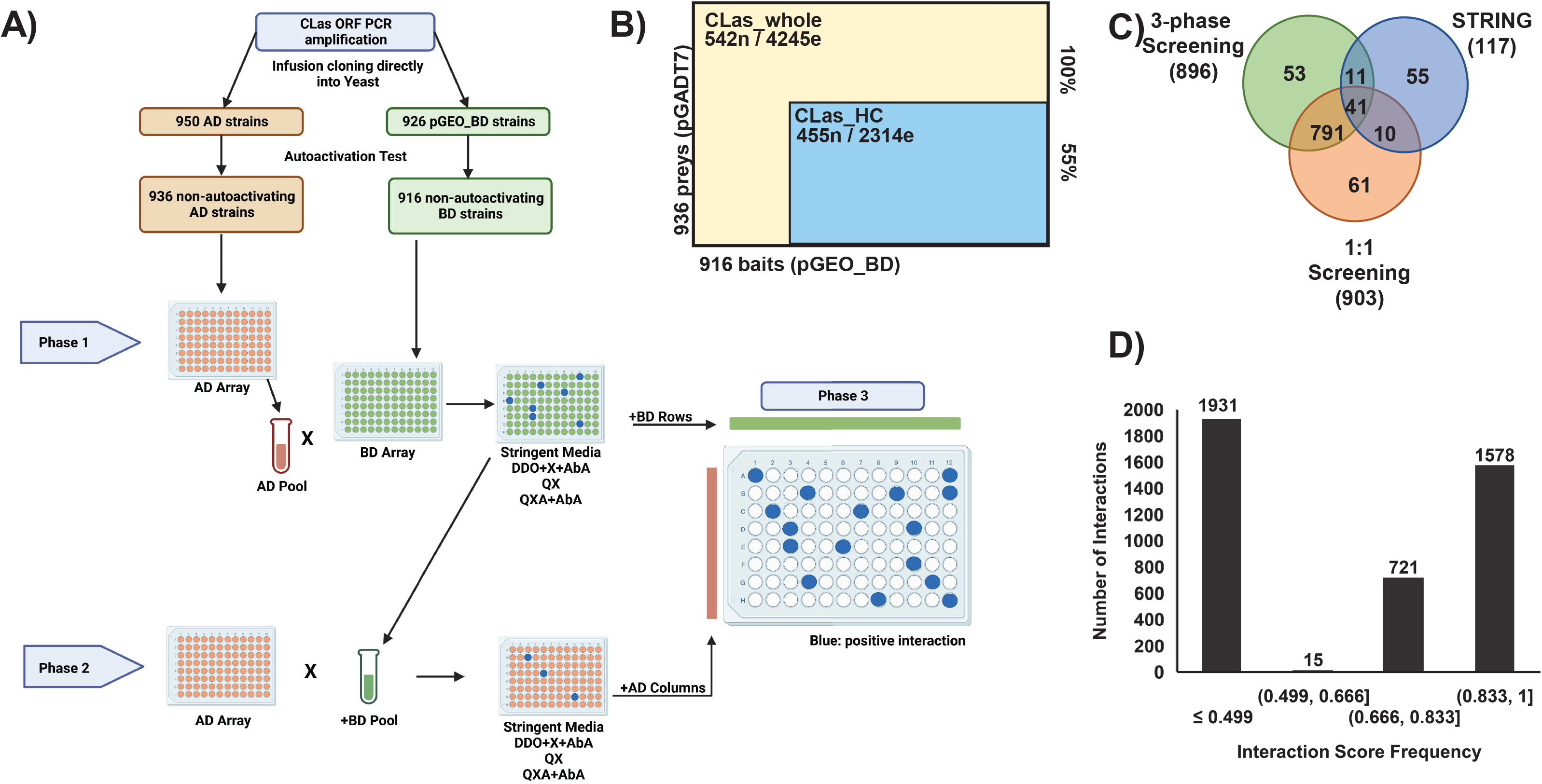
The CLas Y2H interactome screening, validation, and confidence scores. **A.** The three-phase 96-well plate screening pipeline for constructing the CLas_whole network. Phase one and two systematically screened pools and individual yeast constructs to identify potential interacting protein partners. All interacting partners were identified in the third phase in a pairwise screening; phase 1 and 2 interacting proteins were screened pairwise against all other phase 1 and 2 interacting proteins to produce the binary data. **B.** CLas Y2H three-phase screening results. n: node; e: edge; 100% indicates: CLas_whole network; 55% indicates: CLas_HC network portion of the CLas_whole network **C.** Venn diagram showing the overlap of interactions detected in the three-phase screening, an independent pairwise screening, and predicted interactions for 163 CLas proteins; predicted interactions were downloaded from the String consortium. **D.** Interaction confidence score frequency across the CLas_whole network. Interactions with a confidence score 20.5 are considered high confidence; denoted as the CLas_HC network.

### CLas interactome coverage and confidence scores

The robustness of the Y2H datasets is imperative for utilizing the PPI information ^8, 23, 32–34^. We reduced false positives by pairwise rescreening and stringent selection of the CLas network in the three-phase screening. To evaluate the sensitivity and reproducibility of the three-phase method, we subjected a random set of 326 yeast constructs (163 proteins in each vector) to an independent pairwise Y2H screening. We observed 832 reproducible interactions between the three-phase and pairwise screening, representing 92% of the interactions detected between those 163 proteins in the CLas_whole network.

We further employed known PPIs in the STRING-db to evaluate the sensitivity of the two screening assays. Limited information is available for CLas in the STRING database, with no experimentally determined interactions reported for CLas specifically. However, there were 474 and 369 experimentally determined interactions between homologous proteins (iPPIs) and interacting protein families interologs (iCOGs) at confidence cutoffs ≥0.4 and ≥0.9, respectively (Extended Data Table S2; Table S4) ^35, 36^. Our results showed that both Y2H screening methods had comparative sensitivity in detecting predicted CLas interactions based on known PPIs of homolog proteins. Each CLas Y2H screening detected 41 of the 117 predicted interactions among these 163 proteins (Figure 1C, Supplementary Dataset S4).

Using the interactions detected in the two Y2H screening methods, we calculated the false positive rate for the high-throughput three-phase Y2H screening as 3.1% and the false negative rate as 47.9%. Thus, our three-phase Y2H is stringent against false positives in identifying putative PPIs of CLas. However, though the high throughput three-phase screening is efficient, it also brings a significant trade-off of a false negative rate of 47.9%, probably resulting from the pooling used in the 1^st^ and 2^nd^ phases. Nevertheless, the low false positive rate rendered our three-phase Y2H data highly robust to analyze the CLas interactome and infer the protein function of hypothetical proteins.

To evaluate the robustness of individual PPIs, we calculated interaction confidence scores for the CLas_whole network PPIs using interologs as the positive reference set. The interologs were obtained by querying the STRING-db for experimentally determined PPIs from *Agrobacterium radiobacter, B. subtilis*, *C. jejuni*, *E. coli, H. pylori, L. crescens*, *M. genitalium*, *M. loti*, *M. pneumoniae, S. cerevisiae, S. meliloti, T. pallidum,* and iCOGs. There were 635 interologs between 208 protein pairs in the CLas_whole network. We used 100 PPIs corresponding to interologs with the highest confidence scores representing the positive reference set for logistic regression analysis to assign interaction confidence scores to CLas_whole PPIs (Supplementary Datasets S5 and S6). In total, 2314 interactions between 455 proteins demonstrated high confidence scores (≥0.5, CLAS_HC), representing 55% of the total interactions detected in the CLas_whole network and 44% of the CLas ORFs (Figures 1B & 1D, Supplementary Dataset S7)

To further verify our Y2H PPIs, we conducted pull-down assays for nine PPIs focusing on flagellar proteins because of the crucial roles of CLas motility in the HLB pathosystem ^37^. Consistent with the Y2H result, nine tested PPIs were confirmed (Figure 2), supporting the high quality of the three-phase Y2H screening.

**Figure 2.**
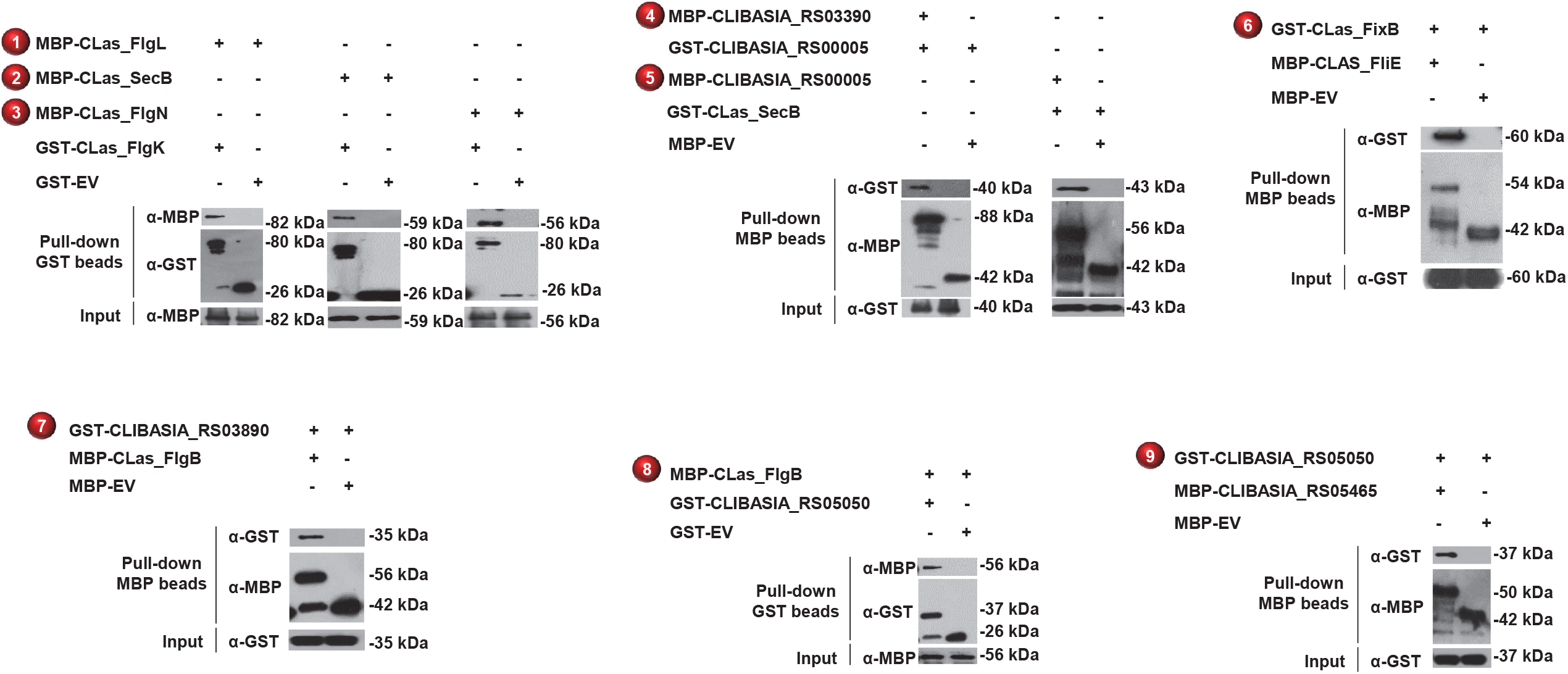
Confirmation of the PPIs identified in Y2H using pull-down assays. Both GST and MBP pull-down assays were conducted, dependent on protein pairs. Interactions confirmed *in vitro* are numbered for illustration in Fig. 5.

### Network Topology

Protein network topology provides valuable information for protein functions^38, 39^. We visualized and analyzed the topology of the CLas Y2H interactome (Figure 3A-C)^40, 41^. The degree distribution topology of CLas_whole had a scale-free organization, typical of a scale-free PPI network (Figure 3D)^27^. The average number of interactions per node was 15.664 in the CLas_whole network; CLas_HC has an average node degree of 10.171. The CLas network had many node neighbors, consistent with the notion that protein function complexity increases with genome reduction (Figure 3F, Extended Data Figure S3) ^6, 42, 43^. The CLas_HC network maintained a degree distribution and topological coefficient distribution like that of the CLas_whole network (Figures 3D and G). However, the clustering coefficient and betweenness centrality distribution of CLas_HC were less dense than the CLas_whole with fewer crosstalk between nodes (Figures 3E and F)^6^.

**Figure 3.**
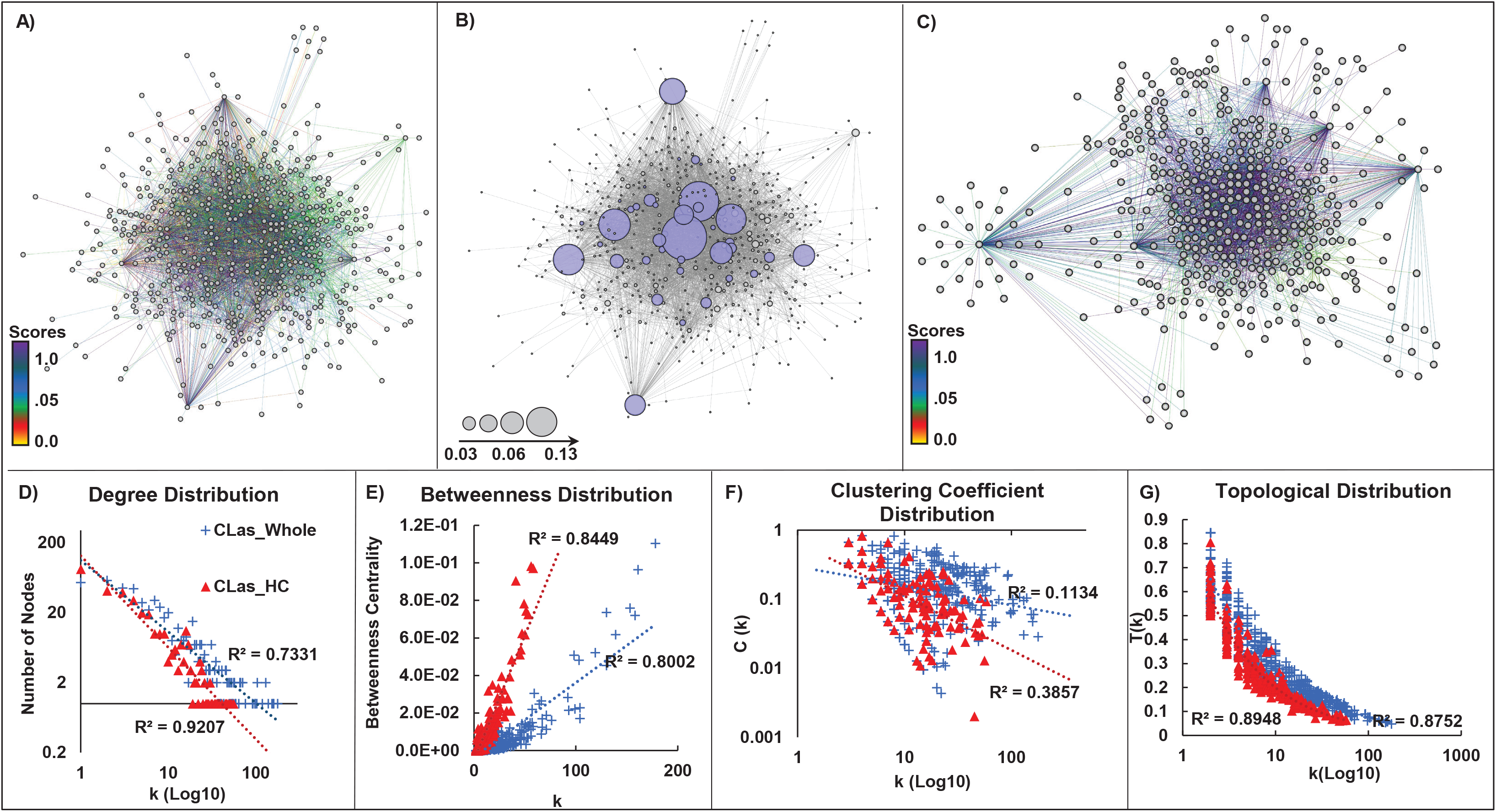
The CLas Y2H network topology. A-C. Graphic representation. n: node indicating a protein in the network; e: edge indicating the connection between the interacting proteins. **A.** CLas_whole network (542n,4245e). **B.** HUB nodes in the CLas_whole network are colored blue; node size is proportional to its betweenness centrality value. **C.** CLas_HC network; PPI confidence scores 20.5. **D-G.** Network topology of the CLas_whole and CLas_HC networks. **D.** Degree distribution. **E.** Betweenness centrality distribution. **F.** Clustering coefficient distribution. **G.** Topological coefficient distribution.

Highly connected nodes (HUBs) are more likely to be responsible for maintaining the overall connectivity of the network. They are more likely to be essential genes or involved in critical PPIs than non-hub nodes ^44^. We identified 40 HUB nodes using manual curation based on network centralities and connectivity, MCC, and MCODE algorithms^44–46^. Using betweenness centrality and degree distribution, we manually curated 27 HUB nodes (Figure 3D-E). We validated the HUBs using the MCODE and MCC clustering algorithms ^45, 46^. MCODE clustering confirmed 26 of our 27 manually curated nodes as HUBs with scores ≥4. The MCC clustering algorithm confirmed 14 HUB nodes defined by manual curation and MCODE clustering and suggested 13 additional HUB proteins (Figure 3B, Supplementary Dataset S8). Interestingly, eight of the HUB proteins were reported to be essential for *L. crescens*^47^, and 24 are known to be essential in other bacteria^48^.

### Inferring functions of CLas hypothetical proteins

CLas contains 415 uncharacterized proteins, including proteins annotated as hypothetical proteins, function unknown, or not in a COG^4, 49^. The CLas_HC network contained 1485 high-confidence interactions involving 160 uncharacterized proteins. The features of these proteins include: 15 have signal peptides that may direct them to the cell membrane or outside the cell; 13 are secreted by a nonclassical pathway that does not involve signal peptides; 27 have transmembrane domains that span the cell membrane; and 107 have no assigned COG that indicates their function^4, 36, 50–52^. One hundred forty-nine uncharacterized proteins interacted with at least one annotated protein, and 105 interacted with two or more proteins with known functions (Supplementary Dataset S3).

To infer the functions of those uncharacterized proteins based on their interacting partners, we employed four guilt-by-association (GBA) methods: 1. Binary interactions and interologs, 2. Operon associations, 3. Protein family associations, and 4. Meta-interactome analysis (Figure 4 illustrates each analysis method)^15, 53–55^.

**Figure 4.**
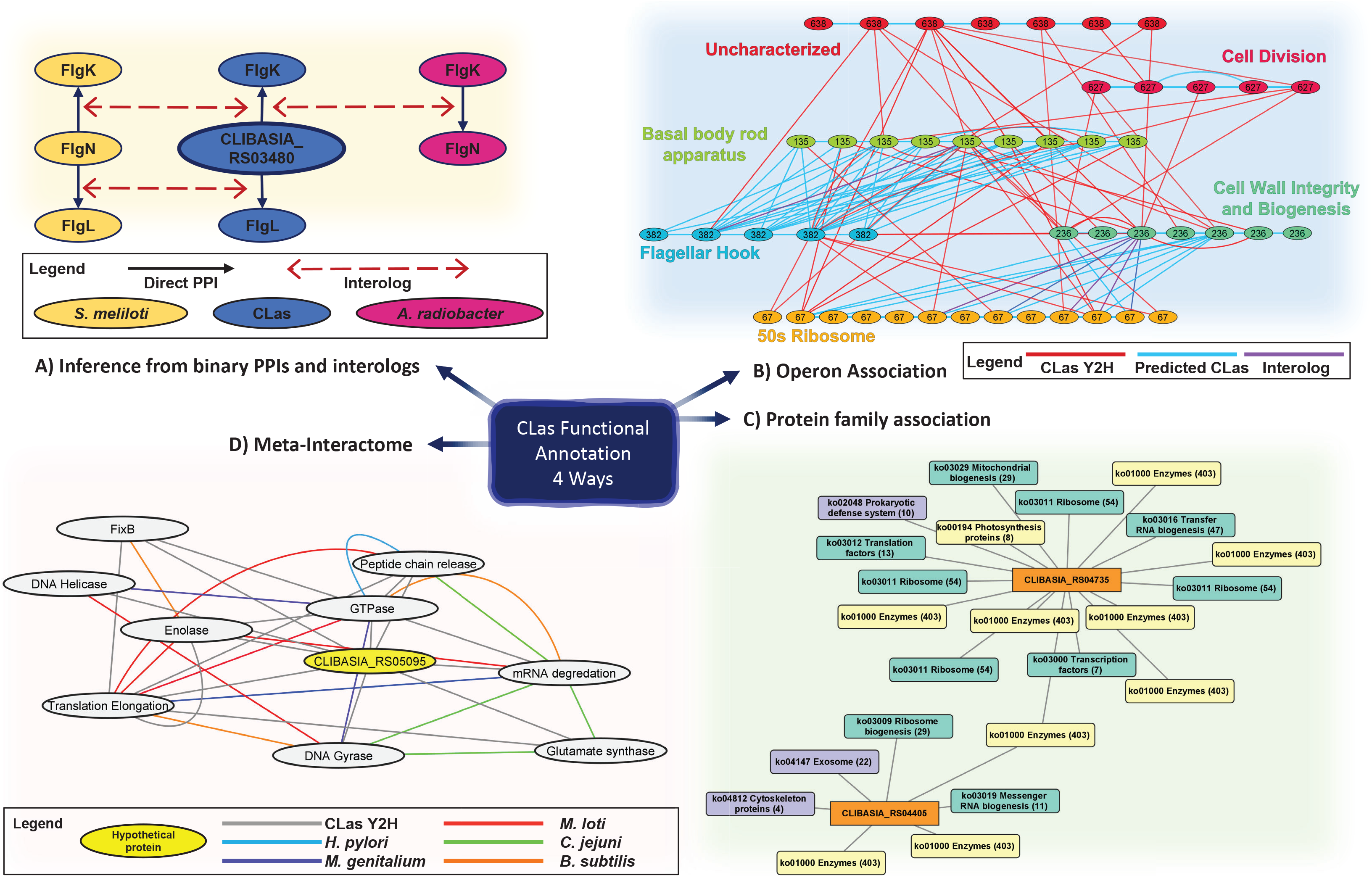
Four approaches were used to infer CLas hypothetical protein fu ctiona notations. **A.** Inter-operon interactions highlight associations between motility related genes, respiration, membrane integrity, ribosomal proteins, and uncharacterized genes and operons. Operon associations, network and properties are in table S9. **B.** Hypothetical protein annotations were implied by interologs and binary interactions. These validity of the interactions were confirmed by *in vitro* assays and tertiary protein structure homology. **C.** Guilt by association was used for CLas proteins interacting with 22 proteins in the same protein family or KEGG pathway. CLIBASIA_RS04405 has a functional association with enzymatic proteins, ribosome and RNA biogenesis, and exosome, cytoskeleton proteins. The binary interactions between CLIBASIA_RS04735 and ribosome, transcription/translation proteins implies a functional association with cell response and growth. **D.** The sub network meta-interactome (mY2H_5095_) for CLIBASIA_RS05095 whose protein function was inferred by its ortholog associations as having a putative association with genetic information processing.

#### Inferring CLas protein functions from binary interactions

Interologs are conserved interactions between proteins whose homologs interact in other organisms ^11^. As aforementioned, we identified 635 interologs between 208 protein pairs in the CLas_whole network (Supplementary Datasets S5 and S6). Among them, 34 were uncharacterized proteins. We have assigned functions to four proteins based on interacting partners and similarities between proteins and their 3D structures (Extended Data Figure S4). CLIBASIA_RS03480 was reannotated as FlgN_Las_, a flagellar switch chaperone, because it interacted with FlgK_Las_ and FlgL_Las_ proteins involved in flagellar hook formation in the CLas network (Figure 4A, Extended Data Figure S4A). The interactions between these proteins are well documented and are interologs found in *S. meliloti* and *A. radiobacter* (Figure 4A)^56^. Similarly, we reannotated CLISASIA_RS03395 as FliK_Las_, a flagellar length regulator, because of its interactions with FlgD_Las_, FlhB_Las_, and FliP_Las_ (Extended Data Figure S4B), three proteins involved in flagellar motor assembly, in the CLas network^57–60^. Additionally, CLas hypothetical proteins CLIBASIA_RS03285 and CLIBASIA_RS03885 were annotated as TolA_Las_ and YcgR_Las_ respectively (Extended Data Figure S4C-D).

#### Inter- and Intra- operon binary interaction topology gives functional association to CLas uncharacterized proteins

Next, we took advantage of the fact that bacterial genes of related functions are transcribed together under a single promoter as operons^61^. We reasoned that operons with known proteins interacting with the operons containing uncharacterized proteins provide functional associations/clues for those groups of uncharacterized proteins. For this purpose, we only considered inter- and intra-operon interactions where two or more proteins from each operon interacted together for further analysis (Supplementary Dataset 9). We examined how the PPIs of other organisms relate to the CLas Y2H network, specifically inter- and intra-operon PPIs (Figure 4B and Extended Data Figure S5). We found that CLas had more inter-operon PPIs than intra-operon interactions (Supplementary Datasets S9 and S10) compared to interactions between homologous proteins. These inter-operon interactions provided evidence of functional association of three CLas operons (638, 603, and 643), which consist of 21 uncharacterized proteins. Specifically, our interactome data demonstrated that CLas operon 638 has a functional association with flagellar assembly, stress response-related proteins, 50s ribosomal proteins, cell division, and cell wall integrity proteins (Figure 4B). In addition, CLas operon 603 has a functional association with translation, FeS cluster assembly, and succinate dehydrogenase. In contrast, CLas operon 643 has a functional association with metabolism, enzymes, and translation, in addition to FeS cluster assembly and succinate dehydrogenase (Extended Data Figure S5).

#### Protein family associations within the CLas Y2H network give clues to protein function

We evaluated PPIs between hypothetical proteins and CLas proteins within known protein families or functions from the Kyoto Encyclopedia of Genes and Genomes (KEGG)^62^. Sixty-one CLas proteins with unknown functions interacted with two or more proteins from the same protein family or function in the CLas_HC Y2H network. These interactors were primarily involved in metabolism and genetic information processing, as shown in examples of protein association networks for CLIBASIA_RS04735 and CLIBASIA_RS04405 (Figure 4C). Operon associations for eleven hypothetical proteins mirrored their KEGG associations, supporting the functional assignment of eleven reannotated proteins (Supplementary Datasets S11 and S12; Extended Data Tables S3 and S4).

#### The CLas “meta-interactome” supports protein family association annotations and provides evidence for additional hypothetical proteins

As a final sweep of the Y2H data, we looked for associations in a meta-interactome^15^. We generated a meta-interactome (mY2H) using experimentally confirmed PPIs for *A. radiobacter, B. subtilis*, *C. jejuni*, *E. coli, H. pylori, L. crescens*, *M. genitalium*, *M. loti*, *M. pneumoniae, S. cerevisiae, S. meliloti,* and *T. pallidum*. The CLas mY2H has 12,780 edges between its 542 proteins (Figures 4D, Extended Data Figure S6). Notably, most interactions (not including the CLas_whole Y2H PPIs) in the mY2H were PPIs in *M. loti*, totaling 2296 edges. *M. loti* is more closely related to CLas than other organisms with available Y2H interactome maps, such as *E. coli* and *H. pylori* ^8, 20, 21, 23^. We identified 416 CLas proteins with orthologs among the 542 proteins in the mY2H, which contained 12,780 PPIs. Of these PPIs, 635 PPI interologs between 208 protein pairs were conserved in the CLas_whole, whereas 4037 were not (Supplementary Dataset S13). We used the orthologous PPI subnetworks and the CLas protein associations from the mY2H to infer the functions of 26 uncharacterized CLas proteins (Figures 4D, Extended Data Figure S6; Supplementary Dataset S12; Extended Data Tables S3 and S4).

### Novel interactions identified for CLas proteins with known functions

The CLas_whole network identified 4245 interactions between 542 proteins (Figure 1B). Among the 371 CLas proteins with known functions (Extended Data Tables S2 and S3), we identified 1654 interactions, whereas their homologs had 171 unique experimentally verified interactions in previous studies ^8, 15–18, 20–23, 63^. This study identified 1483 novel PPIs that have not been previously reported for proteins with known functions (Supplementary Dataset S14).

### Y2H PPIs related to Flagellar proteins

Flagellar motility might play critical roles in CLas infection of different organs of psyllids and movement in phloem tissues. Importantly, active movement of CLas was observed in the phloem tissue against the flow of phloem sap^37, 64^. We paid close attention to flagella-related proteins. Our Y2H screening detected a total of 399 interactions among known flagellar proteins in our Y2H screening (Fig. 5; Supplementary Dataset S15; Extended Data Tables S6, S7, and S8). In addition, nine interactions were confirmed with pull-down assays (Fig. 2). Of those PPIs confirmed, two were putative secreted proteins, CLIBASIA_RS05050^52^ and CLIBASIA_RS05465^50^, interacting with each other and FlgB_Las_, a flagellar basal-body protein^65^. Another secreted protein, CLIBASIA_RS03890, also interacted with FlgB_Las_^65^. These interactions imply that these proteins may play a role in flagellar formation and functions, which could affect the pathogenicity or survival of CLas^50, 66–68^. FlgN is a known flagellar export chaperone of FlgK and FlgL, two hook-filament junction proteins^56^. We also identified SecB, a chaperone protein dedicated to translocating proteins across the cytoplasmic membrane, as an interactor of FlgK. We validated both interactions, FlgK_Las_-FlgN_Las_ and FlgK_Las_-SecB_Las_, consistent with the involvement of the Sec pathway in flagellar synthesis^69, 70^. Moreover, SecB_Las_ interacted with CLIBASIA_RS0005, a protein with T-SNARE-like domains. T-SNARE proteins mediate membrane fusion in eukaryotic cells. Pathogens can exploit host SNARE proteins to enter host cells by displaying SNARE-like domains on their surface membrane^71^. It remains to be determined whether CLIBASIA_RS0005 is involved in CLas entering cells or glands of psyllids^72^.

**Figure 5.**
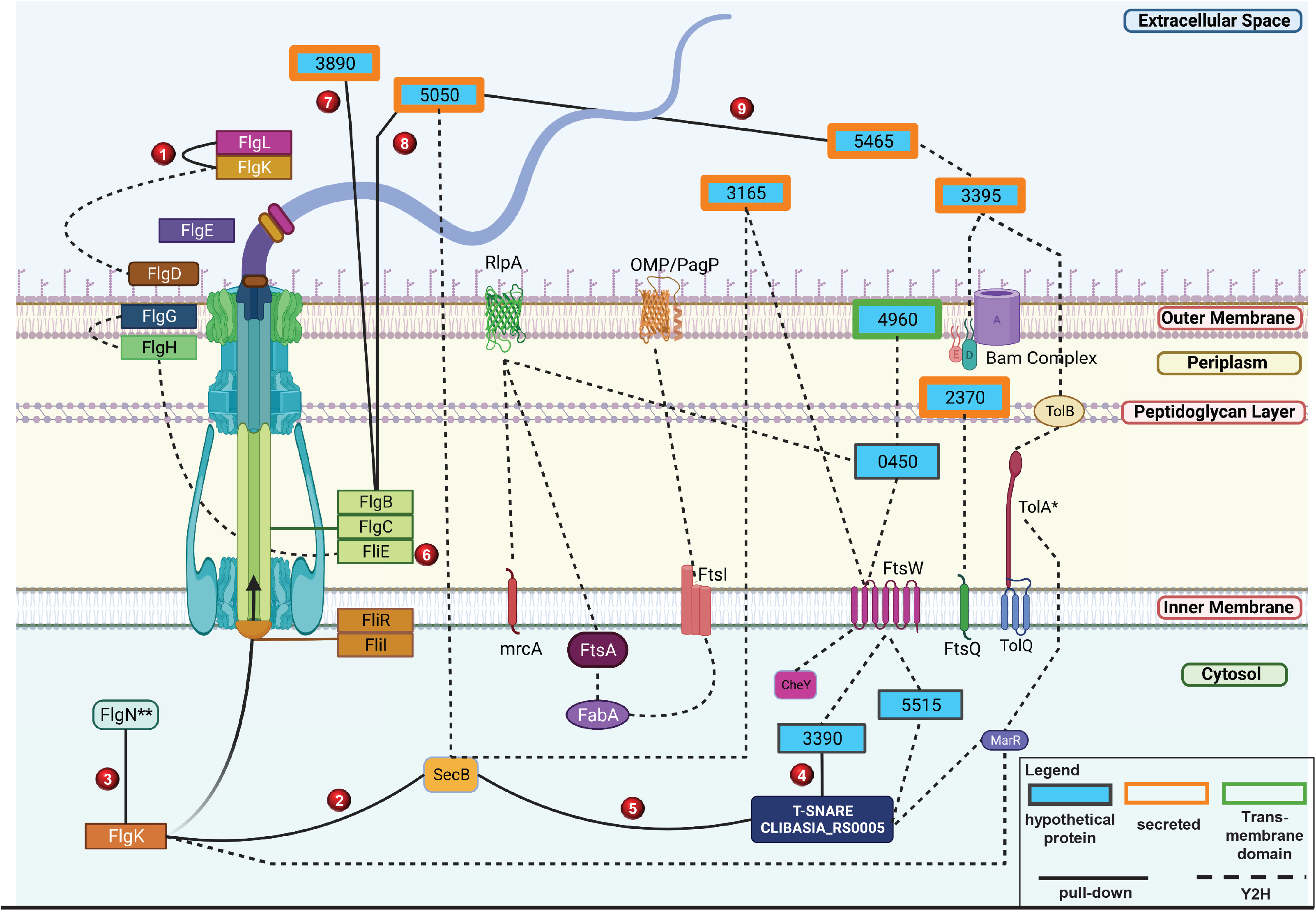
Integrated view of CLas proteins related to flagellar biosynthesis and membrane proteins, their interacting partners identified in Y2H. Subcellular localization representation of CLas hypothetical and annotated protein interactions. The source of the interactions and hypothetical proteins are denoted by dashed and solid lines. Hypothetical proteins with orange borders are experimentally confirmed secreted proteins as described in Prasad et al. (2016) and Du et al. (2021). Interactions confirmed with pull-down experiments were numbered from 1-9 in red circles. The hypothetical proteins were named using their CLIBASIA_RS#.

## Discussion

We have constructed the first and only large-scale protein-protein interaction map of CLas using a high-throughput Y2H approach. In total, we have identified 4245 PPIs among CLas proteins. We have employed a three-phase screening approach to improve the robustness of the Y2H. We estimated that our high-throughput Y2H data has a false positive rate of 3.1%, consistent with previously reported false positive rates of 0.5%^73^, less than 5%^74^, and 6.5%^75^. The pull-down assay supported the robustness of our Y2H.

Two hundred and eight binary PPIs, of the 4245 PPIs detected in this study, had been reported in previous studies^7, 8, 15–23, 35, 63, 76^. On the other hand, 4037 novel interactions were identified in this study, expanding the PPI database. Two thousand five hundred eighty-three interactions involved 171 uncharacterized CLas proteins were identified, providing clues for their functions. For instance, we have assigned protein functions to CLIBASIA_RS03480 as FlgN, CLIBASIA_RS01260 as FliK, CLIBASIA_RS03285 as TolA, and CLIBASIA_RS03885 as YcgR. CLIBASIA_RS04960 and CLIBASIA_RS02370 might play a role in cell division since they interact with FtsW_Las_ and FtsQ_Las_, two essential components of the divisome complex^77^.

These proteins interact with a HUB protein CLIBASIA_RS00450, CLIBASIA_RS05515, and RlpA (septal ring lytic transglycosylase, CLIBASIA_4130; RlpA_Las_). RlpA is known to be involved in cell separation and rod shape of bacteria^78^. Thus, even though the functions of CLIBASIA_RS04960, CLIBASIA_RS02370, CLIBASIA_RS00450, and CLIBASIA_RS05515 are unknown, they are probably involved in cell division or cell wall formation, providing hints for further characterization. We have identified 37 uncharacterized proteins that interact with multiple proteins of an operon, and 61 uncharacterized proteins interacted with two or more proteins from the same family or function, providing strong indications regarding their roles associated with the known functions of the operons or protein families compared to single interactions. Among the 1662 interactions between proteins with known functions of CLas, only 171 have been observed in other systems, indicating that CLas proteins might have new functions to adapt to the reduced genome size. CLas, *T. pallidum*, and *M. pneumoniae* have a greater number of interactions per node in their PPI networks than free-living bacteria with larger genomes, such as *E. coli*, *M. loti*, *H. pylori*, *C. jejuni*, and *B. subtilis*. An increase in node neighbors seems to be a common phenomenon for bacteria that have undergone reductive evolution^42^.

We identified 40 putative HUB nodes in the network ^45, 79–81^. HUB proteins are suggested to be essential to the organism or involved in critical PPIs^22, 38, 79, 80, 82^. Eight of the HUB proteins were reported to be essential for *L. crescens*^47^, and 24 are known to be essential in other bacteria^48^. Interestingly, multiple HUB proteins of CLas protect the bacteria from oxidative stress damage, including quinone oxidoreductase 1^83^ and LysR-type transcriptional regulator^84^. Quinone oxidoreductase 1 acts as a cytoplasmic antioxidant, whereas LysR is a key circuit component in regulating microbial stress responses and is required for bacterial tolerance to H_2_O_2_ in vitro^84^.

The protective mechanism of CLas is consistent with the model that HLB is a pathogen-triggered immune disease; CLas triggers systemic and chronic overproduction of reactive oxygen species, leading to phloem tissue cell death and HLB symptoms^85^. Thus, those proteins will protect CLas from the high ROS levels in the phloem tissues. Another HUB protein CtrA, is known to regulate bacterial cells, such as *Caulobacter crescentus*, to generate cells of different morphologies: with and without flagella^86^. Interestingly, CLas seems to encode most flagellar genes even though flagella have not been observed in CLas^37^. The extensive PPIs involving flagellar proteins suggest they are functional, consistent with the active movement of CLas against the flow of phloem sap ^37, 64^. CLas was also reported to be polymorphic in shape^37^. CtrA may be involved in the non-flagellated and polymorphic phenotypes of CLas cells in phloem tissues. Furthermore, ten of the 40 HUB genes are significantly up-or down-regulated in either citrus or ACP vector^87^, suggesting they play crucial roles for CLas survival in citrus or psyllids.

Overall, we have generated the interactome map for CLas, which has provided insights regarding the biology and pathogenicity of CLas and the putative functions of uncharacterized proteins.

This interactome will serve as a valuable resource and provide clues to further characterize the functions of unknown proteins. The PPIs identified have the potential to be used as targets for the development of novel antimicrobials to control HLB.

## Methods

### Yeast Two-hybrid vector construction

Full-length CLas ORFs were PCR amplified using genomic DNA extracted from CLas-positive citrus leaf tissues. The gel-purified amplicons were fused with the activation (AD) and binding domain (BD) Y2H expression vectors (pGADT7 and pGEO_BD, respectively) using infusion recombinase cloning (Clontech). We modified the Gal4 Matchmaker Gold Yeast Two-hybrid system (clonetech) pGBK_T7 (Clontech) expression vector to harbor the pGADT7_AD vector’s multiple cloning site (MCS) between RE sites NdeI and BamHI (Extended Data Figure S1B) and was named pGEO_BD. This modification simplified the cloning of all CLas ORFs into both vectors between RE sites EcoRI and XmaI using one infusion primer set per gene to reduce costs and labor. These AD and BD infusion reactions (∼20 µl per 50 µl reaction) were directly transformed into yeast strains Y187 (AD, prey clones; MATa; G1::lacZ, M1::MEL1) and Y2H Gold (BD, bait clones; MATα; G1::HIS3, G2::ADE2, M1::AUR1-C, M1::MEL1) following Yeastmaker Yeast Transformation protocol (Clontech). We verified transformants with colony PCR and sequencing a random set of ∼200 constructs.

### Autoactivation test

Autoactivation of the reporter genes by individual ORFs was assessed by mating the transformants with the opposite corresponding mating partner empty vector and plating on selective media DDO/X/AbA (-Leu, -Trp, + XαGal, +Aureobasidin A). Clones that autonomously activated reporter genes were removed from the 96-well plates and high throughput screening.

### Yeast Two-hybrid three-phase screening

A three-phase 96-well mating scheme was adapted from a matrix approach^32^. Briefly, all transformants were cultured individually in 96-well plates in 1 ml of the appropriate minimal liquid media (AD: -Leu; BD: -Trp) at 30°C until saturated (∼36-48 hours). For the first mating phase (three phases in total), all individual 1 ml AD cultures were pooled together and centrifuged at 700 g for 5 min. The pellet was resuspended in 25 ml SD/-Leu liquid media with 25% glycerol. Aliquots (1 ml) of the pool were stored at -80°C. The BD cultures were left in a 96-well format for the phase 1 mating.

### Mating procedure

***Phase 1***. Each AD pool was mated to individual pGEO_BD clones in a 96- well format. The matings were spotted on YPDA agar plates by spotting 3 μl of the AD pool aliquot and 3 μl of the individual BD clone on top of one another and incubated at 30°C for ∼ 48 hrs, or until the colonies were ∼ 1 mm in diameter. The 1 mm colonies were transferred using a pin replicator to 96-well plates with DDO (-Leu, -Trp) liquid medium to select diploids harboring AD and BD plasmids and incubated at 30°C and 210 rpm on a rotary shaker. This step reduces the background on the selection plates. After 48 hrs, five μl of the mating cultures were plated on DDO (-Leu, -Trp) agar plates as a mating control, and selective media to determine interaction and interaction intensity: D/X/A (-Leu, -Trp, + XαGal, +Aureobasidin A), Q/X (-Leu, -Trp, -His, -Ade + XαGal), and Q/X/A (-Leu, -Trp, -His, -Ade + XαGal, +Aureobasidin A). All subsequent matings and screenings were carried out in the same manner. ***Phase 2.*** Positive BD clones from the phase 1 mating were pooled using the previously described AD pooling method. These BD pools (containing all positive BD interactors from phase 1) were mated back to the AD plate that comprised the pool of phase 1. This is to determine which of the AD clones within the pool were the interacting partners to the positive BD interactors from phase 1 (Figure 1A). ***Phase 3.*** The final phase determined the interacting partners from the previous two screenings and allowed for a pairwise rescreening of all interacting constructs. All positive AD and BD yeast constructs interacting in the first two phase screenings were organized into a row (BD) and column (AD) format in 96-well plates. Each row of a 96-well plate is a single BD construct (For example, row A: BD construct #1, row B: BD construct #2, row C: BD construct #3… to row H: BD construct #8) and each column is a single AD construct (For example column 1: AD construct #1, column 2: AD construct #2, column 3: AD construct #3… to column 12: AD construct #12) which were mated pairwise. The row and column format allowed all phase one and two proteins to interact to determine the interacting partners and the interaction specificity. This method helped determine what interactions are more likely, or accurate since interactions must be repeatable pairwise and allowed us to weed out sticky interactors.

### Analysis of the CLas Y2H network topology

We visualized, analyzed, and utilized the network developed in this study with Cytoscape network software (Shannon et al., 2003). Network topologies, such as betweenness and closeness centrality, degree, and clustering coefficient, were determined using the Network Analyzer in Cytoscape (Assenov et al., 2008). We used this information to select HUB nodes in the network for further analysis and understand general CLas interactome network organization found in the Y2H screening. To validate the CLas_whole network topology, NetworkRandomizer version 3 was used to compare the CLas_whole network to a randomly computed network. We compared the difference between the real network (CLas_whole) and its most similar random network for each centrality^88^.

Hub nodes were determined using manual curation, MCODE clustering, and MCC algorithm^44, 89, 90^.

### Scoring interactions and assigning interaction confidence values

PPIs were scored on a scale from one to three for each selection media. These colony scores were used with PPI interologs, PPIs within the same operon, and bait out-degree and prey in-degree to create an additive score for the logistic regression formula to assign interaction confidence scores. The logistic regression training set of 100 interologs was used as true positives and 100 PPIs with the lowest combined Y2H score and highest average shortest path link value with randomized bait and prey node degree values, as true negatives. We calculated the interaction scoring four times using different variables for each regression. The variables were: the number of times as a bait, number of times as a prey, interolog score alone, and interolog score combined with Y2H repeatability and interaction intensity. We averaged the values from each regression to obtain the final interaction confidence score ^16^.

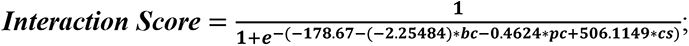

*bc: count of occurrence as a bait node; pc: count of occurrence of prey node; cs: colony score*

### Conserved protein-protein interactions (interologs)

CLas homologs in *A. radiobacter, B. subtilis*, *C. jejuni*, *E. coli, H. pylori, L. crescens*, *M. genitalium*, *M. loti*, *M. pneumoniae, S. cerevisiae, S. meliloti, T. pallidum,* and iCOGs were collected using IMG/MER and NCBI databases ^16, 21, 91, 92^. Only protein sequences with an identity of >30% were considered for analysis. All available experimentally determined (i.e., Y2H, AP/MS, and CO-IP) ortholog PPIs and iCOG PPIs were collected from the STRING database for these homologs^16^. The known and predicted interactions between CLas proteins were also included in the data collected from STRING^35^.

### Validating CLas interactions *in vivo* by a pairwise Y2H screening

We randomly chose 163 ORF constructs (15.9% of the total CLas ORFs and 30% of the nodes in the CLas_whole network) to use in a pairwise Y2H screening. The interactions from this screening were compared for overlap between the three datasets: the pairwise screening, the three-phase Y2H PPI screening, and the known and predicted interactions between these proteins available from the STRING database. The rates of false positives, false negatives, specificity, and sensitivity within the three-phase screening were determined using the known interactions and pairwise screening^93^.

### Confirming CLas interactions using pull-down assay

Interactions were validated using MBP/GST pull-down assay. The pGEX-4T-1 (GST) (GE Healthcare) and pMAL-c5x (MBP) (NEB) vectors were used to construct the *E. coli* expression vectors for GST-Glutathione and/or MBP-Amylose pull-down assay between multiple CLas proteins. The GST/MBP expression vectors were constructed by amplifying the ORFs from the CLas yeast expression vectors and inserted between RE sites: BamHI and SalI in pGEX vector and XmnI and EcoRI in pMAL vector using infusion cloning (Clontech). These newly generated vector constructs were transformed into *E. coli* Rosetta (DE3) electrocompetent cells, transformants were selected by plating on LB agar plates containing ampicillin (100 µg/ml) and chloramphenicol (30 µg/ml); all subsequent steps used these antibiotics at these concentrations. The transformants were cultured overnight in 3 ml liquid LB. The next day, the culture was diluted 1:100 in 25 ml LB and grown until the culture reached an OD_600_ 0.2-0.3. The culture was then induced with IPTG (1 µg/ml) overnight at 16°C.

After induction, the resulting cell pellet was lysed using 5 µg/ml of pellet with B-PER™ Bacterial Protein Extraction Reagent (Thermo-Fisher) and incubated at room temperature for 20 minutes while rocking. The lysates were cleared by centrifugation at 4°C for 20 minutes at 16,000 g. Supernatants were collected in a fresh tube and total protein concentration was measured using Bradford assay. The protein expression and solubility were evaluated by both Coomassie stain and western blot (WB) of the GST and MBP fusion proteins using anti-GST (abcam) and anti-MBP (NEB) monoclonal antibodies before proceeding. The GST-fusion protein and empty-GST supernatants were mixed with the MBP-fusion protein and incubated with glutathione or amylose agarose beads for 3-4 hours at 4°C with rotating. The glutathione or amylose agarose was then washed 5-10 times to remove unbound proteins before boiling and analysis by SDS-PAGE WB using anti-MBP (NEB) and anti-GST (abcam) monoclonal antibodies. Protein-protein interactions that were further confirmed by pull-down assays were shown in T Extended Data Table S9.

## Data availability

The data related to Y2H assays were included in the datasets or supplementary tables.

## Acknowledgements

We thank Wang lab members for constructive suggestions and insightful discussions. This project was supported by funding from Florida Citrus Initiative Program, Citrus Research and Development Foundation, U.S. Department of Agriculture National Institute of Food and Agriculture grants 2022-70029-38471, 2021-67013-34588, 2018-70016-27412 and 2016-70016-24833, FDACS Specialty Crop Block Grant Program, and Hatch project [FLA-CRC-005979] (N. Wang).

## Author Contributions

N.W. and E.W.C. conceptualized and designed the experiments. E.W.C. and O.G.P. designed the pGEO vector and oligonucleotides. E.W.C. performed the experiments. E.W.C. and N.W. wrote the manuscript with input from all co-authors.

## Competing interests

All other authors declare no competing financial interests.

**Correspondence and requests** for materials should be addressed to N. Wang.

---

**Extended Data Table S1.** Previous bacterial interactome studies

| Organism | Type of Organism | Total ORFs in Genome | Proteins Interacting in Y2H | % ORF Covered | Total Interactions | Avg. Number PPI per Node | System | Reference |
| --- | --- | --- | --- | --- | --- | --- | --- | --- |
| *M. loti* | gram - | 7277 | 1804 | 24.8 | 3121 | 1.7 | ‡Gal4 | Shimoda et al.<br>(DNA research, 2008) |
| *B. subtilis* | gram + | 4100 | 287 | 7 | 793 | 2.8 | ‡Gal4 | Marchadier et al.<br>(Proteomics, 2011) |
| *Synechocystis sp.* | gram - | 3692 | 1920 | 52 | 3236 | 1.7 | Gal4 | Sato et al.<br>(DNA research, 2007) |
| *E. coli* | gram - | 3600 | 1269 | 35.3 | 2234 | 1.8 | Gal4 | Uetz et al.<br>(Nature biotechnology, 2014) |
| *S. pneumoniae* | gram + | 2109 | 820 | 38.9 | 2045 | 2.5 | LexA | Wuchtly et al.<br>(ASM molecular biology and physiology, 2017) |
| *C. jejuni* | gram - | 1654 | 1321 | 79.9 | 2884 | 2.2 | LexA | Parrish et al. (Genome Biology, 2007) |
| 1st *H. pylori* | gram - | 1587 | 261 | 16.4 | 908 | 3.5 | Gal4 | Rain et al.<br>(Nature, 2001) |
| 2nd *H. pylori* | gram - | 1587 | 739 | 46.6 | 1515 | 2.1 | Gal4 | Hausert et al.<br>(Molecular and Cellular Proteomics, 2014) |
| “Comprehensive”<br>*H. pylori* | gram - | 1587 | 1135 | 71.5 | 2369 | 2.1 | Gal4 | Rain & Hausert et al.<br>(Combined Study) |
| *T. pallidum* * | gram - | 1039 | 726 | 69.9 | 3649 | 5 | Gal4 | Titz et al.<br>(PLOS One, 2008) |
| CLas* | gram - | 1027 | 542 | 52.8 | 4245 | 7.8 | Gal4 | This Study |
| *M. pneumoniae* | gram + | 689 | 617 | 90 | 10083 | 16.34 | TAP-MS | Kuhner et al. (Science, 2009) |

\*Obligate intracellular; ‡: Not a complete interactome.

**Extended Data Table S2.**
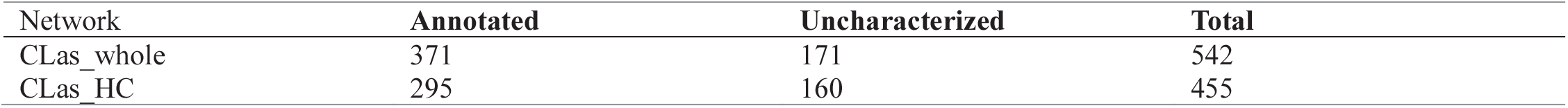
Number of annotated and uncharacterized proteins in the CLas_whole and CLas_HC networks

| Network | Annotated | Uncharacterized | Total |
| --- | --- | --- | --- |
| Clas_whole | 371 | 171 | 542 |
| Clas_HC | 295 | 160 | 455 |

**Extended Data Table S3.** Interactions between annotated and uncharacterized proteins.

| Interaction type | Clas_whole<br>Node | Clas_whole edge | Clas_HC<br>node | Clas_HC edge |
| --- | --- | --- | --- | --- |
| Annotated - annotated | 339 | 1491 | 246 | 692 |
| Annotated - hypothetical | 502 | 1976 | 388 | 1106 |
| Hypothetical - hypothetical | 148 | 570 | 123 | 388 |
| Interologs (annotated – annotated) | 128 | 171 | 96 | 106 |
| Interologs (annotated – hypothetical) | 47 | 30 | 30 | 18 |
| Interologs (hypothetical - hypothetical) | 12 | 7 | 7 | 4 |
| Total | 542 | 4245 | 455 | 2314 |

**Extended Data Table S4.** 542 CLas proteins were queried for their predicted interactions available from the STRING-db.

|  | STRING-db 0.4<sup>a</sup> |  | STRING-db 0.9<sup>b</sup> |  |
| --- | --- | --- | --- | --- |
|  | nodes | edges | nodes | edges |
| All STRING association types | 436 | 3509 | 256 | 863 |
| Experimentally determined | 115 | 474 | 81 | 369 |

Note: Two STRING datasets of experimentally proven PPIs (interaction confidence scores of $\geq 0.4^a$ and $\geq 0.9^b$ ). The resulting network node and edge counts are listed in the table. No lab experiment datasets determined for Clas are available from STRING. All interaction associations are predicted based on experiments conducted in orthologs.

**Extended Data Table S5.** Count of uncharacterized proteins by each annotation evidence/method used.

| Annotation Method | Number of uncharacterized proteins |
| --- | --- |
| protein families' association | 39 |
| metainteractome | 16 |
| operon association | 12 |
| protein families' association; operon association | 11 |
| PPI | 7 |
| metainteractome; protein families association | 6 |
| PPI; structure | 4 |
| metainteractome; operon association | 2 |
| protein families' association; PPI | 1 |
| protein families' association; structure | 1 |
| metainteractome; PPI | 1 |
| metainteractome; protein families association; PPI | 1 |
| Total | 101 |

**Extended Data Table S6.** PPIs between 22 flagellar proteins in the CLas Y2H.

| Interaction type | Clas_whole<br>Node | Clas_whole edge | Clas_HC<br>node | Clas_HC edge |
| --- | --- | --- | --- | --- |
| annotated Flg proteins – all PPIs | 213 | 399 | 118 | 175 |
| annotated Flg proteins – annotated protein | 122 | 240 | 59 | 103 |
| annotated Flg proteins – uncharacterized | 69 | 151 | 40 | 67 |
| annotated Flg proteins – annotated Flg<br>proteins | 9 | 8 | 7 | 5 |

**Extended Data Table S7.**
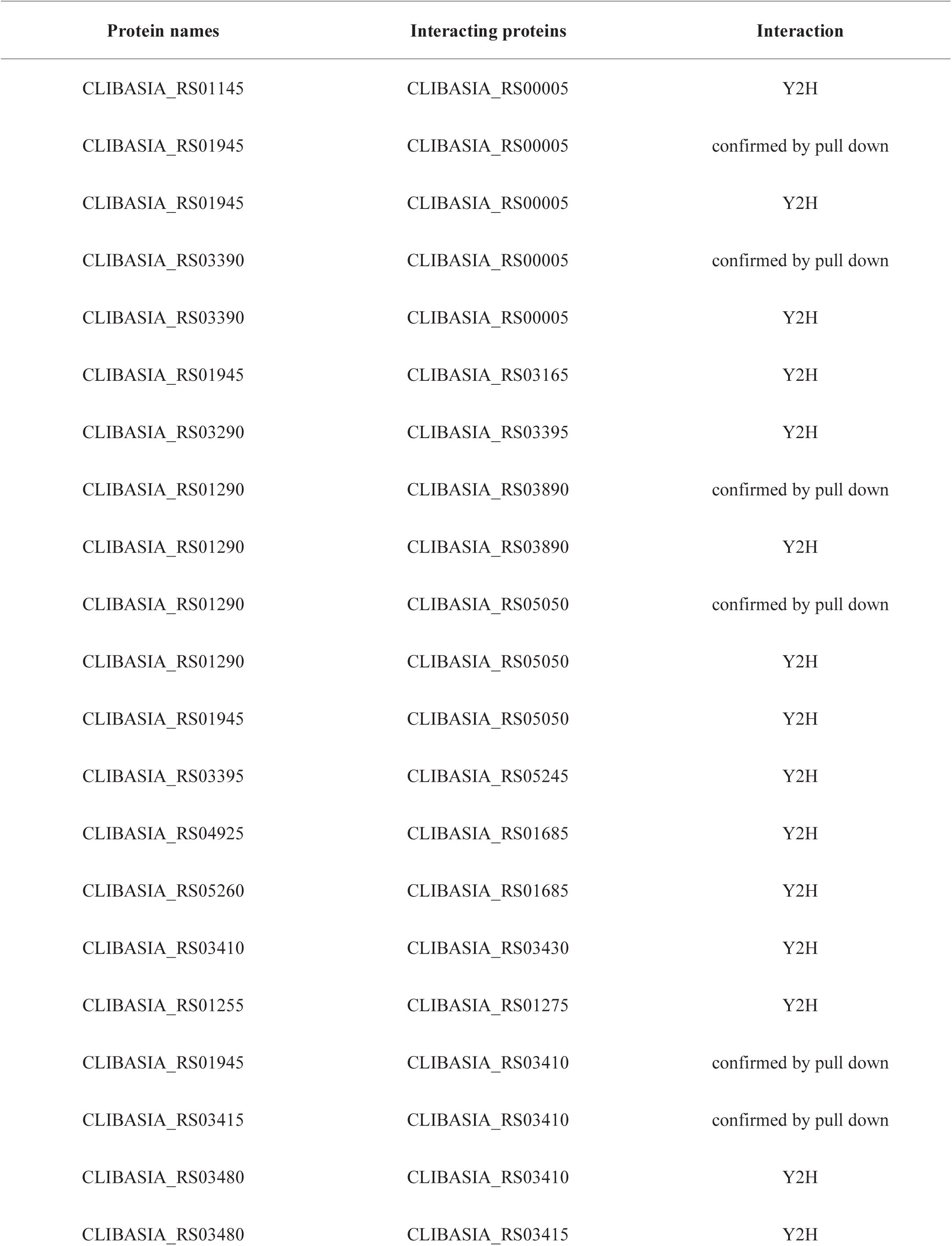
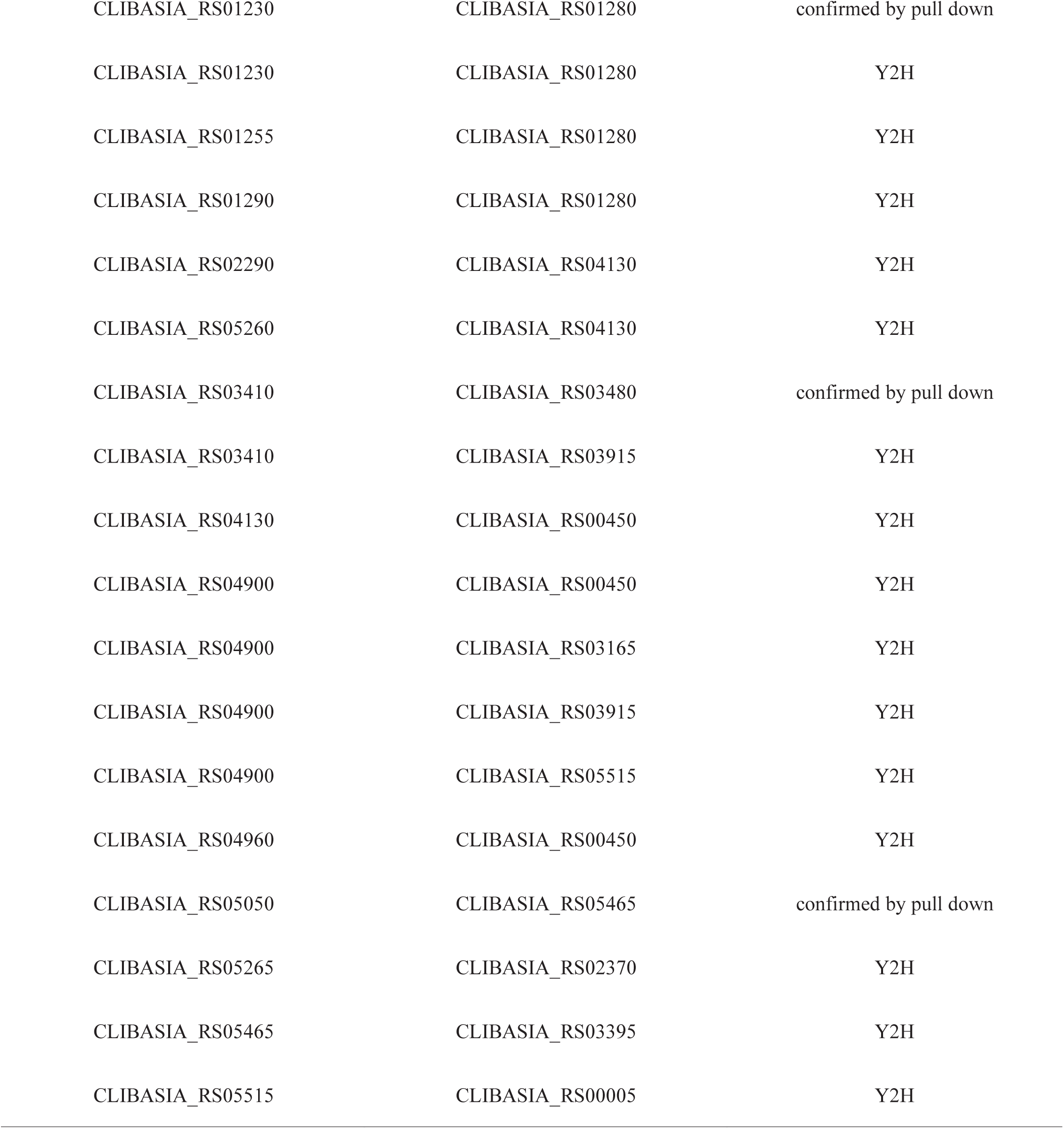
CLas PPIs involving flagellar and membrane proteins.

**Extended Data Table S8.**
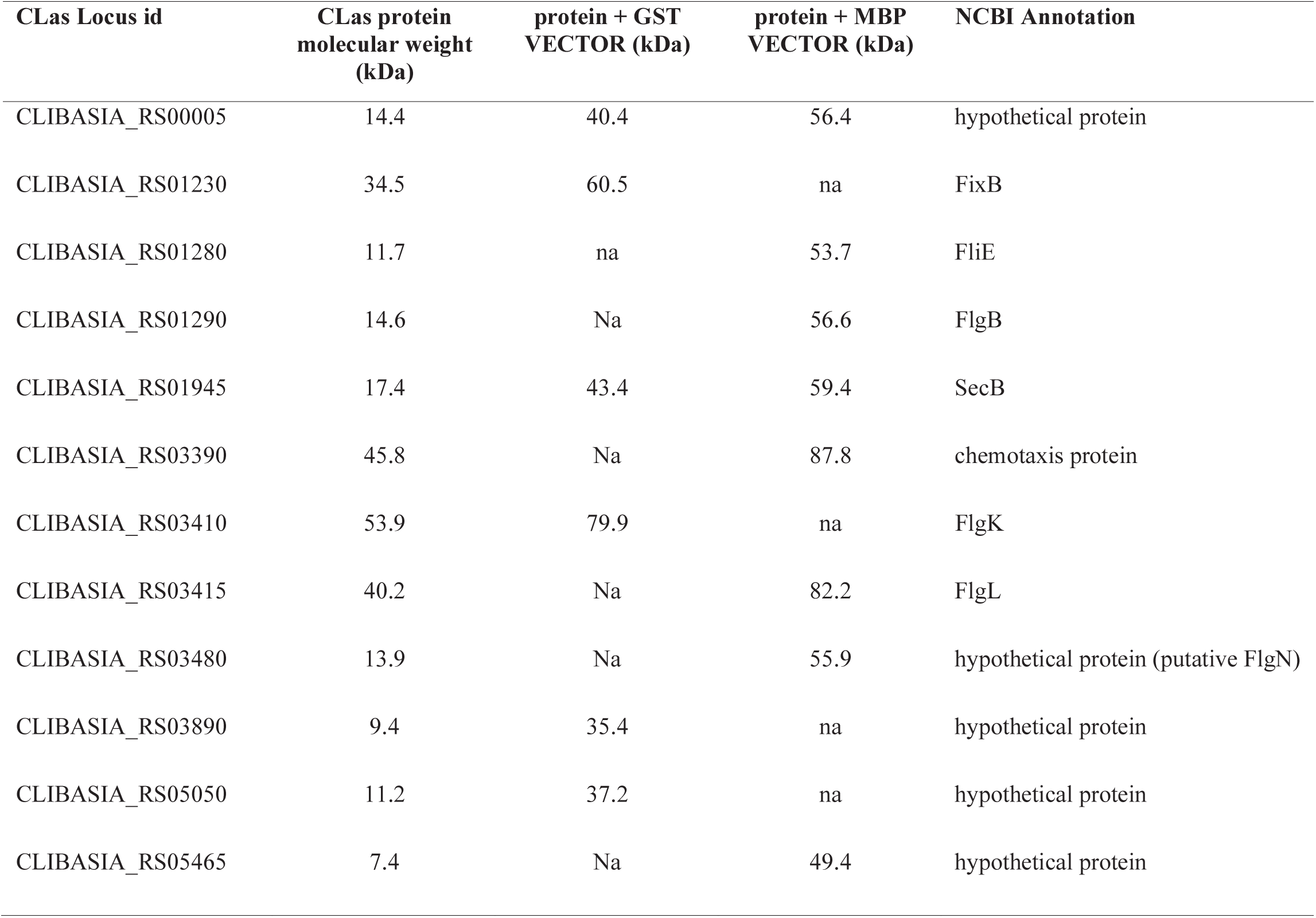
CLas protein-protein interactions that were confirmed by pull-down assay

**Extended Data Figure S1.**
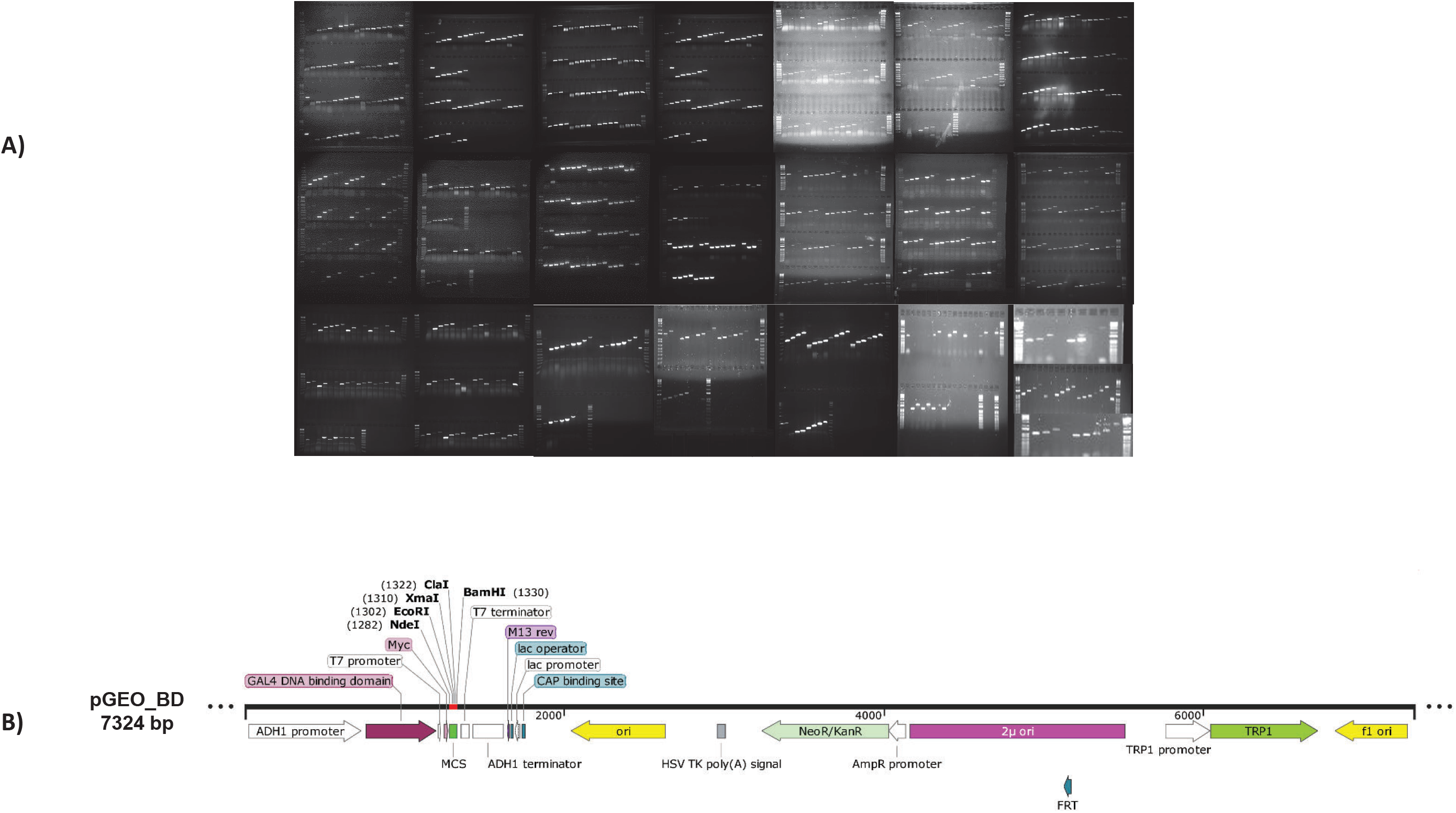
Cloning of CLas genes for Y2H. A) Amplification of CLas ORFs for infusion cloning. Primers for 1027 CLas ORFs were designed, resulting in 974 successfully amplified genes for Y2H high throughput interaction screening. **B) Modification of Yeast Two-Hybrid expression vectors enables efficient cloning of all CLas ORFs into two vectors.** A fragment containing the pGADT7 AD (clonetech) MCS sequence between NdeI and BamHI was used to replace the MCS in pGBKT7 (clonetech) between the restriction sites NdeI and BamHI to create a new binding domain vector pGEO_BD. The modified MCS allowed us to design only one primer set for each CLas gene to clone into both AD and BD vectors using Infusion cloning (Clonetech) for the Y2H high-throughput screening. MCS: multiple cloning site.

**Extended Data Figure S2.**
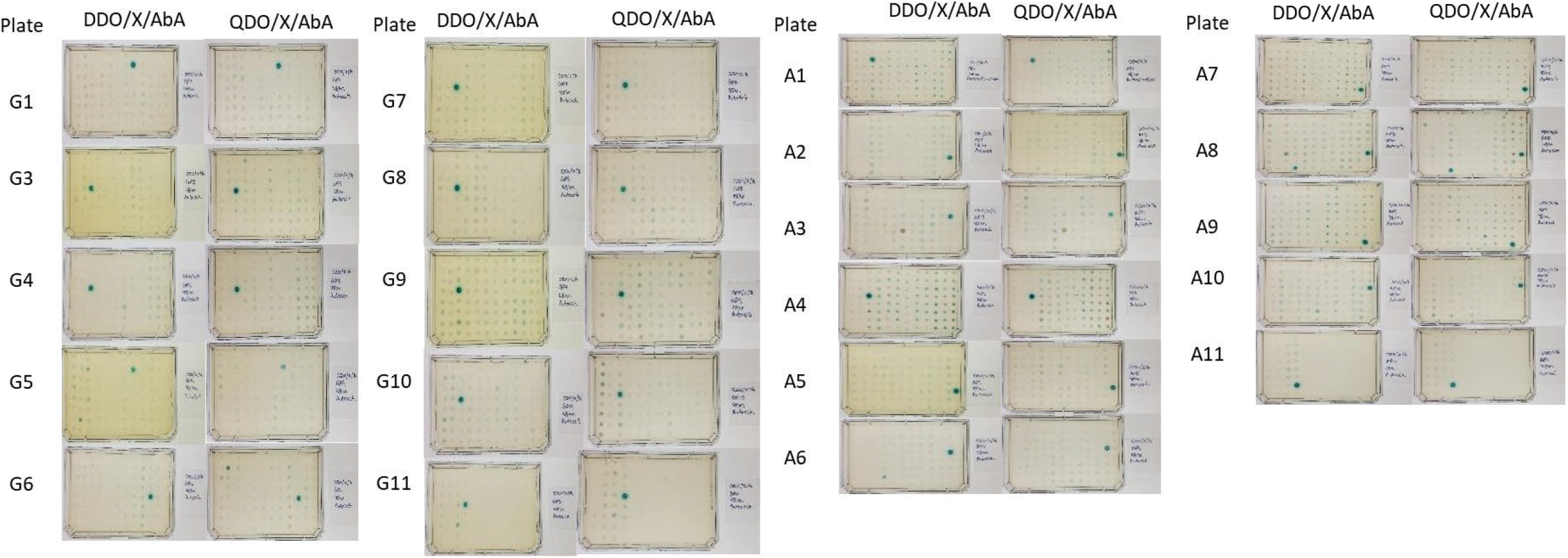
Constructing and testing yeast expression vectors. Autoactivation screening for CLas ORF constructs in both AD and BD vectors. From the ORFs cloned into both the AD and BD Y2H vectors, constructs were transformed into Mat-a yeast and mated with Mat-a yeast harboring the empty vector opposite of the construct; i.e., AD construct mated with BD empty vector, vice versa.

**Extended Data Figure S3.**
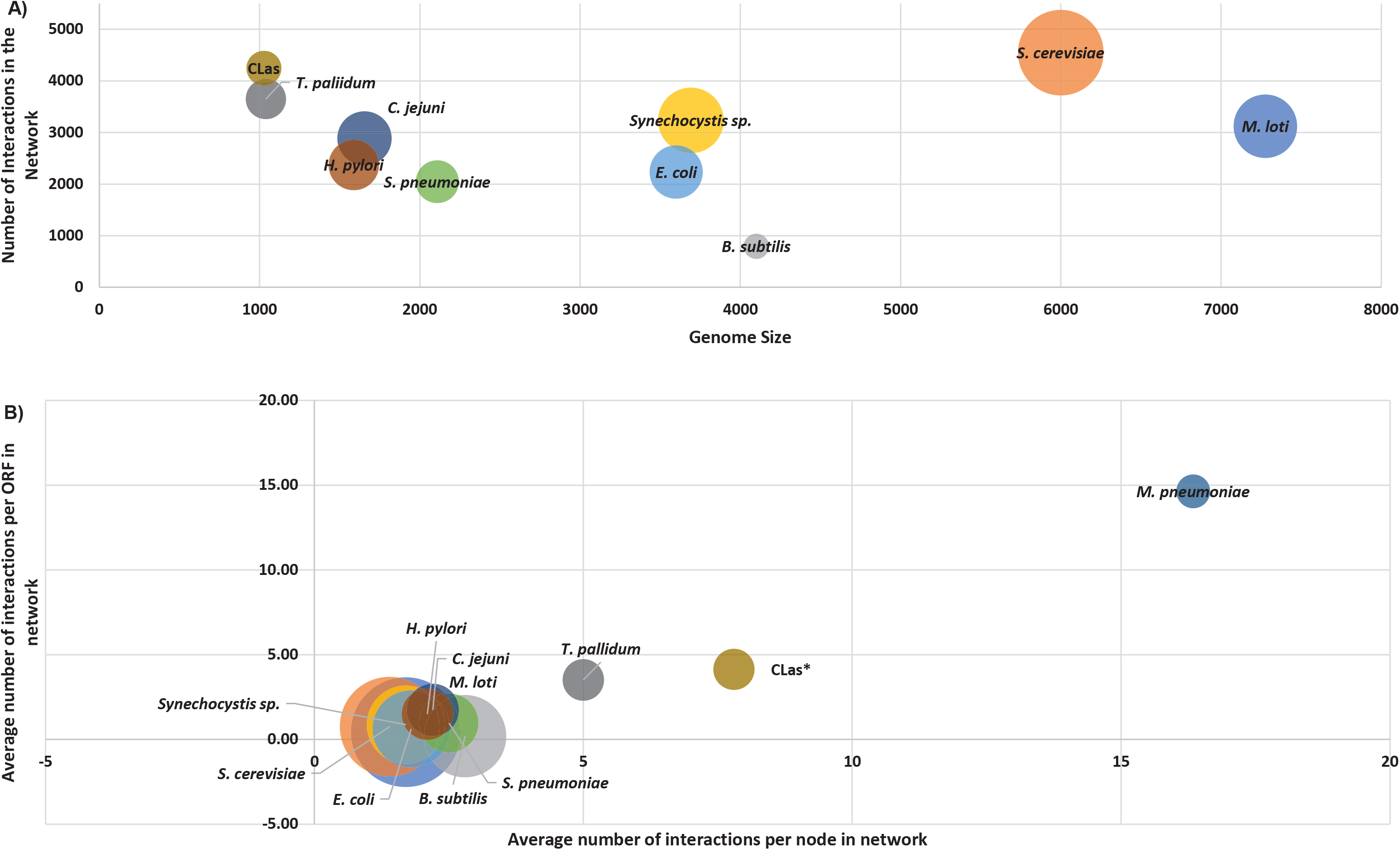
Comparison of CLas interactome with other bacteria. **A.** Two interactomes of bacteria with reduced genome sizes (CLas and *T. pallidum*) have a higher number of interactions between fewer number of proteins, compared to other interactomes. Here, the genome size, number of nodes, and number of interactions were compared across nine interactomes. Bubble size represents number of nodes in a network. **B.** Relationship between genome size and the average number of interactions per node. The CLas_whole is shown in comparison to other interactomes. Bubble size represents genome size.

**Extended Data Figure S4.**
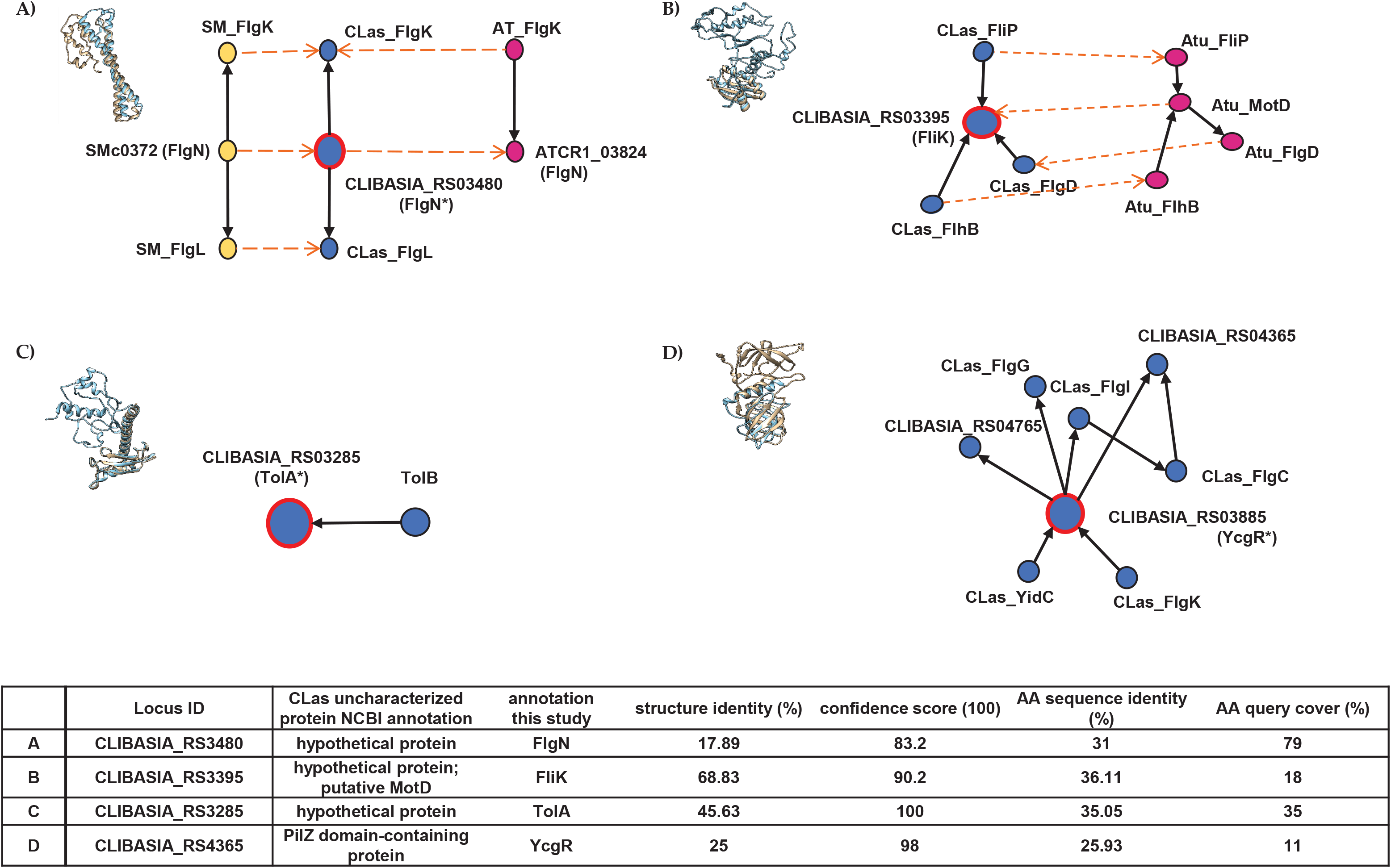
Protein-protein interactions, interologs, and protein structure homology data were used to assign putative functions for CLas uncharacterized proteins. The Phyre2 online database was used to predict CLas protein structures and find possible template matches of HMM homolog proteins and characterized proteins. Protein structure alignments of CLas proteins (blue), aligned and superimposed over protein Phyre2 protein templates (gold). Alignment and confidence data for each are shown below the figures**. A)** Protein structure alignment of CLIBASIA_RS03480 and *Bradythizobium* FlgN. The interologs in *S. meliloti* (yellow) and *A. tumefaciens* (purple) suggest that CLIBASIA_RS03480 To be FlgN. **B)** CLIBASIA_RS3395 was annotated to be FliK based on structure alignment and interologs in *A. bacterium*. **C)** CLIBASIA_RS3285 was annotated to be TolA based on structure alignment and its interaction with CLas TolB in the Y2H network. **D)** CLIBASIA_RS4365 was annotated to be YcgR flagellar brake protein based on structure alignment and interacting partners in Y2H. * annotation was conducted in this study.

**Extended Data Figure S5.**
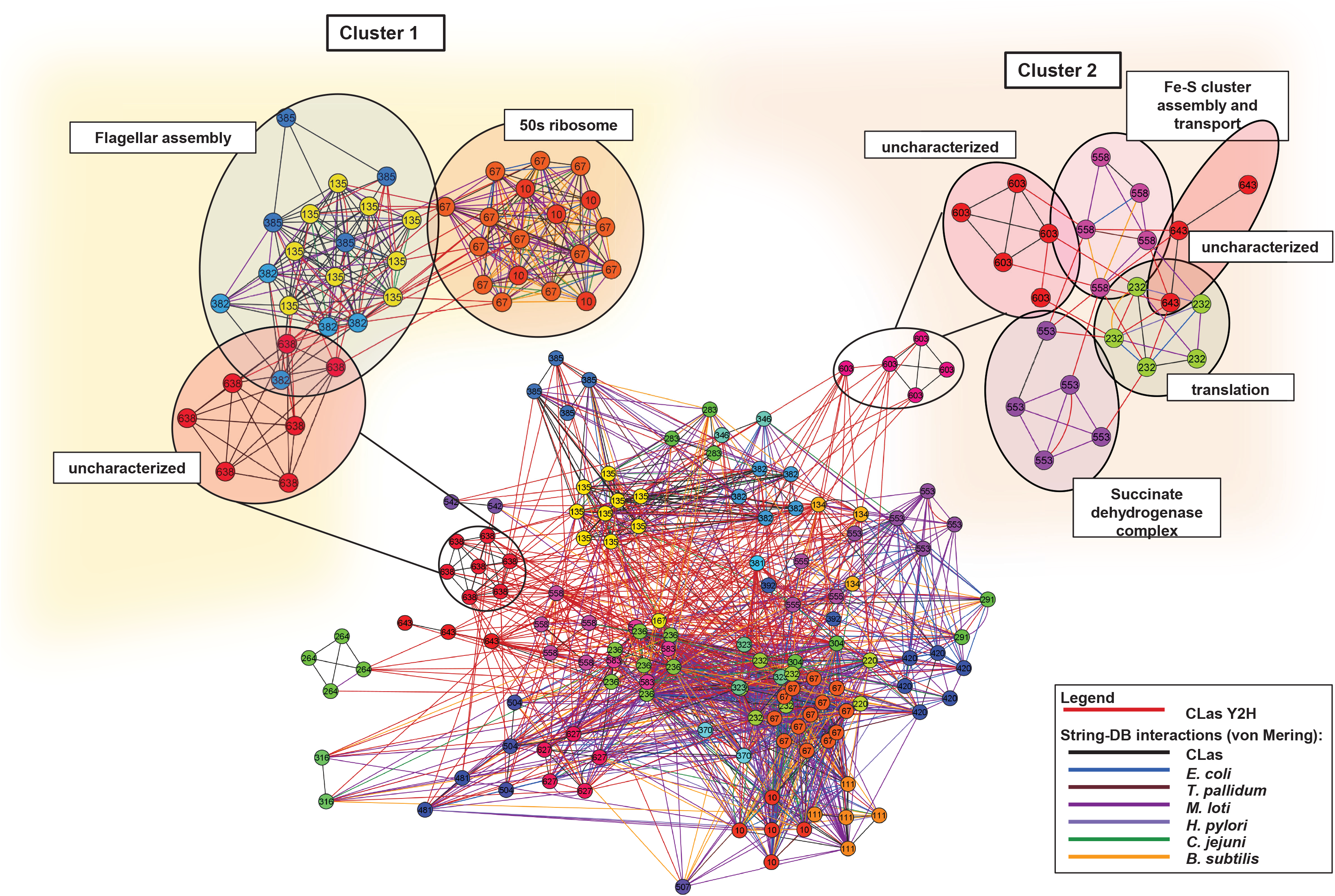
Inter-operon interactions highlight associations between motility related genes, respiration, membrane integrity, ribosomal proteins, and uncharacterized genes and operons. The large network shown below the two subnetwork clusters is the CLas operon network. Operons were predicted using the ProOpDB: Prokaryotic Operon Data Base (http://operons.ibt.unam.mx/OperonPredictor). CLas operons with 22 PPIs with another operon are shown. Three operonic gene clusters of uncharacterized proteins interact with annotated CLas proteins; uncharacterized operons are in red; i.e.: 603, 638, and 643. Edges are colored by PPI source. The CLas Y2H PPIs in this study are represented by red edges.

**Extended Data Figure S6.**
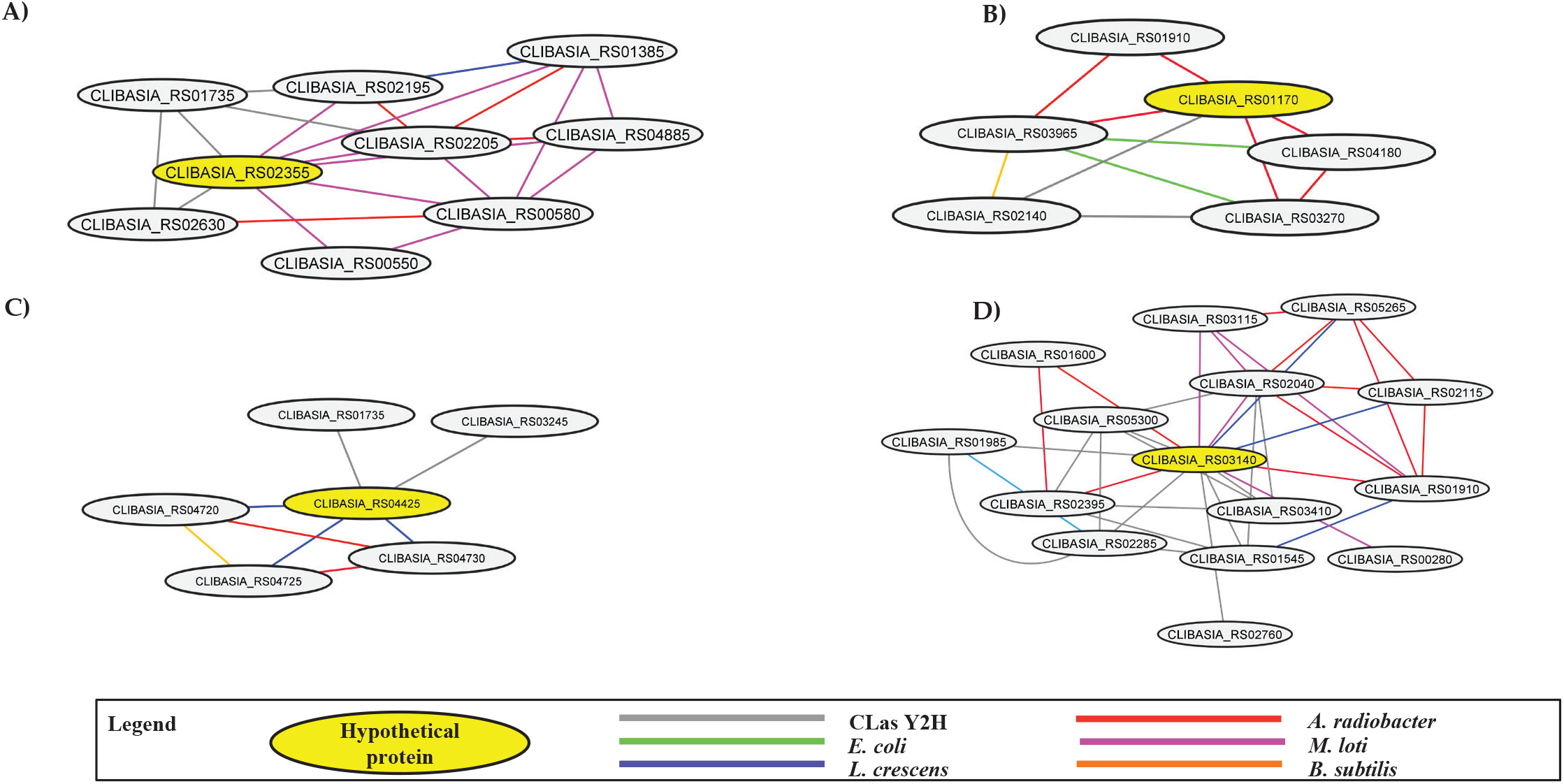
Meta-interactome subnetworks. Four hypothetical proteins whose protein function was inferred by its ortholog associations found in meta-interactome (mY2H) subnetworks. Edges are colored by ortholog PPI. A. CLIBASIA_RS02355 has a putative association with cell wall formation and lipid synthesis. B. CLIBASIA_RS04425 has a putative association with FeS cluster assembly, DNA YacG, and RNA pyro phosphohydrolase. C. CLIBASIA_RS0170 has a putative association with DNA processing related proteins. D. CLIBASIA_RS03140 has a putative association with cell division.

